# Dysregulated TGFβ-ERK Signaling Drives Aberrant Extracellular Matrix Production in Noonan Syndrome-Associated Pulmonary Valve Stenosis

**DOI:** 10.64898/2026.01.16.700032

**Authors:** Clifford Z. Liu, Shrey Patel, Simone Sidoli, Aditi Prasad, Daniel Charytonowicz, Amine Mazine, Alexander A. Mikryukov, Jamshid Abdul-Ghafar, Elizabeth S. Kahn, Dejauwne Young, George A. Porter, Philip J. Katzman, Stephen P. Sanders, Chrystalle Katte Carreon, Dirk Hubmacher, Joy Lincoln, Gordon M. Keller, Wendy K. Chung, Robert Sebra, Bruce D. Gelb

## Abstract

Pulmonary valve stenosis (PVS) is the most common congenital heart defect in Noonan syndrome (NS) and related RASopathies, yet the molecular mechanisms linking pathogenic variants to the valve pathology remain poorly defined. Here, we utilized a human iPSC-based valve differentiation platform to generate the cardiac valve cell lineages—including fibrosa and spongiosa valve interstitial cell (VIC) subtypes. CRISPR-edited iPSCs harboring NS gain-of-function RAS/MAPK and Noonan syndrome with multiple lentigines (NSML) dominant-negative RAS/MAPK variants exhibited early defects in mesodermal and endocardial specification in all genotypes. Additionally, NS-iPSC endocardial cells exhibited defects in endothelial-to-mesenchymal transition (EndMT) specifically towards fibrosa VICs, which was most pronounced in *PTPN11^N308D^* (N308D) cells. Single-cell transcriptomics revealed widespread dysregulation of extracellular matrix (ECM) programs in N308D fibrosa VICs, including increased expression of collagens and proteoglycans, as well as dysregulation of multiple genes involved in ECM remodeling. We also detected activation of RAS-MAPK, TGFβ, and fibrosis-associated pathways in our transcriptional dataset. Mass spectrometry-based phosphoproteomics confirmed coordinated increases in ERK, PKC, and stress-related kinases, as well as enhanced activity of the TGFβ receptor. Functionally, N308D fibrosa VICs exhibited exaggerated upregulation of ECM genes in the presence of TGFβ2 ligand, suggesting that these cells are hypersensitive to TGFβ stimulation. Furthermore, we demonstrated that this pathological ECM-program occurs independently of *BAMBI*, a negative regulator of TGFβ signaling that was found to be decreased in N308D fibrosa VICs. Lastly, we performed histopathological analyses of stenotic pulmonary valves from two NS infants, which demonstrated marked overproduction and disorganization of ECM, mirroring the findings from our iPSC-based disease model. Together, our data reveal a central mechanism where NS-associated alleles sensitize fibrosa VICs to TGFβ, which leads to aberrant downstream signaling and drives the pathological ECM program in NS-associated PVS.

## Introduction

Noonan syndrome (NS) and related RASopathies arise from pathogenic germline variants that perturb signaling through the RAS-MAPK pathway. Two related RASopathies, NS and Noonan syndrome with multiple lentigines (NSML; formerly LEOPARD syndrome), share significant overlapping clinical features, yet their underlying variants have distinct biochemical effects. NS alleles typically enhance RAS-ERK signaling, while NSML alleles act in a dominant-negative manner and instead augment PI3K-AKT-mTOR signaling.^1,2^ These disorders present with a broad spectrum of clinical findings, but cardiac involvement is especially prominent and ranges from hypertrophic cardiomyopathy to congenital heart disease.^3,4^ Across the spectrum of congenital heart disease, pulmonary valve stenosis (PVS) is the most common lesion, affecting approximately half of individuals with NS and 10-20% of those with NSML. Involvement of the other heart valves, though much rarer, can also occur in these conditions.^5–7^ Pathological variants in *PTPN11*, which encodes for the tyrosine phosphatase SHP-2 and accounts for about half of NS and the vast majority of NSML cases, has a particularly strong association with PVS, occurring in over 80% of individuals.^8,9^ In contrast, *RAF1* variants rarely lead to valve disease in NS, instead causing severe hypertrophic cardiomyopathy in the majority of patients.^10,11^

During normal human development, valvulogenesis begins when endocardial cells overlaying the cardiac jelly undergo tightly regulated endothelial-to-mesenchymal transition (EndMT), giving rise to nascent valvular interstitial cells (VICs) that migrate into the cardiac jelly and populate the endocardial cushions.^12,13^ Subsequent VIC proliferation and extracellular matrix (ECM) deposition drive cushion growth and elongation to form primitive valve leaflets. Throughout fetal development and into childhood, the valve leaflet ECM undergoes extensive remodeling, ultimately yielding the mature trilaminar structure of the adult valve: a collagen-rich fibrosa layer, a proteoglycan- and glycosaminoglycan-rich spongiosa layer, and an elastin-rich ventricularis/atrialis layer.^14–17^ Proper formation and maintenance of these layers are essential for normal valve function, as disruptions in ECM remodeling or homeostasis are hallmarks of valve pathology.^16,18^

Despite the high prevalence of PVS in NS and NSML patients, the underlying molecular mechanisms remain unclear. Traditional mouse models of NS have provided important insights into the early developmental changes associated with the valve phenotype in NS but also have significant limitations. In these mice, the valvular phenotype is incompletely penetrant, with adult survivors typically lacking valve disease and, when cardiac defects are present, mid-gestational embryonic lethality occurs, limiting investigations to fetal tissues.^19,20^ As such, mechanistic investigation has relied on endocardial cushion explants from mouse and chick embryos. One such study identified enlarged endocardial cushions by E13.5 in NS *Ptpn11^Q79R^*mice and identified an ERK1-dependent increase in proliferation and reduction in apoptosis within the cushion mesenchyme.^21^ Similarly, enhanced ERK signaling and prolonged EndMT were shown to drive increased production and migration of mesenchymal cells in *Ptpn11^D61G^* NS mouse cushion explants.^20^ When chick cushion explants were infected with adenoviruses expressing the *PTPN11^Q79R^* allele, an ERK-dependent increase in proliferation was observed in cushion mesenchymal cells.^22^ Intriguingly, NSML *Ptpn11^Q510E^* mouse explants did not demonstrate increased proliferation but instead exhibited increased cell migration.^23^ Beside these studies, there has been a paucity of mechanistic investigation into this pathology. Notably, there have been no investigations using human valvular cells and histological analyses of diseased pulmonary valves from patients with NS have not, to the best of our knowledge, been described. This absence of human mechanistic and tissue-level data leaves a major gap in the understanding of how RASopathy variants shape human valve development and disease.

To address these challenges, our group recently developed a feeder-free human iPSC differentiation system that produces VICs with fibrosa*-* or spongiosa*-*like identities.^24,25^ In the present study, we applied this system to CRISPR-engineered iPSCs carrying NS (*RAF1^S257L^*, *PTPN11^N308D^*) or NSML (*PTPN11^Y279C^*) variants in order to define their effects on human valve development and function. Through phenotypic characterization during differentiation, we identified key developmental stages that are aberrant in our iPSCs harboring pathogenic variants. Combining transcriptomic and phosphoproteomic analyses, we detected pronounced abnormalities in ECM production, accompanied by dysregulation of TGFβ and RAS-MAPK signaling pathways, in *PTPN11^N308D^* fibrosa VICs. *In vitro* experiments suggested that this excessive production of ECM reflects an exaggerated sensitivity to TGFβ, rather than differences in TGFβ ligand levels. Finally, histopathological analysis of stenotic pulmonary valves from two infants with NS demonstrated dramatic overproduction and disorganization of ECM, mirroring the phenotypes observed in our *in vitro* iPSC-derived model.

Together, these findings provide evidence for a mechanistic link between NS variants, the dysregulation of ERK and TGFβ signaling, and the pathological production and remodeling of ECM in the human pulmonary valve. Importantly, outcomes from this study serve as a foundation for understanding the molecular mechanisms that drive NS-associated PVS and enable the future development of targeted therapies for these patients.

## Methods

Detailed descriptions of cell culture, generation of iPSC cell lines, differentiation protocols, flow cytometry, single-cell RNA sequencing, mass spectrometry-based phosphoproteomics, and histopathological analysis of pulmonary valve tissues are available in the **Supplemental Methods**.

### Institutional Approval

Studies involving the use of human pulmonary valve tissue were conducted in collaboration with Boston Children’s Hospital and The Stella and Richard Van Praagh Cardiac Registry (Institutional Review Board (IRB) #P00046264) and Columbia University (IRB #AAAA5720). All samples were de-identified and collected with the signed consent of parents in accordance with all relevant institutional, state, and federal regulations. Only information pertaining to patient age and clinical diagnoses were collected.

### Materials and data availability

Data generated from single-cell RNA sequencing were uploaded to Gene Expression Omnibus (GEO) and can be accessed with the accession number GSE315927. Data generated from the mass spectrometry phosphoproteomic experiments are available on the public repository ProteomeXchange (PRIDE) under the Project Accession PXD071910. This study did not generate any original code. Any additional information or materials needed to support the findings from this study are available from the corresponding author upon reasonable request.

### Quantification and statistical analysis

All data are represented as mean ± standard error of mean (SEM). Biological replicates of each differentiation experiment are represented by sample size (N). Sample sizes were not predetermined and due to the nature of the experiments, samples were not randomized or blinded from investigators. Statistical significance was determined with either one-way ANOVA, two-way ANOVA, or the Student’s t test. Multiple comparisons with Sidak, Tukey, or Dunnet’s post hoc test were performed as needed in GraphPad Prism 10 software. Results were marked with significance cutoffs of p < 0.05 (*), p < 0.01 (**), p < 0.001 (***), and p < 0.0001 (****). In instances where adjusted p-values were reported, these values are Bonferroni corrected. Biological effect sizes for scRNAseq transcriptomics were determined using Cohen’s *d* and classified as small (0.5 < |*d*| < 1.0), moderate (1.0 < |*d*| < 1.5), or large (|*d*| > 1.5). All statistical tests and parameters are reported in the respective figures and figure legends.

## Results

### Generation of NS and NSML iPSC lines

We began our studies with the well-characterized WTC11 human iPSC line^26^ and used CRISPR/Cas9 genome editing to introduce NS- and NSML-associated variants into this common genetic background. Specifically, we generated iPSC lines harboring NS variants that confer gain-of-function activity on the RAS-MAPK pathway, including *RAF1^S257L/+^* (S257L)—a variant strongly associated with severe hypertrophic cardiomyopathy and negatively associated with PVS—as well as *PTPN11^N308D/+^* (N308D^Het^), the most common NS variant and one that is strongly associated with PVS.^9,10^

Although patients with the *PTPN11* N308D variant are almost invariably heterozygous, prior work by Araki *et al.* demonstrated that mice heterozygous for this allele do not exhibit cardiac defects, whereas homozygous embryos display either severe, lethal cardiac abnormalities that include enlarged valve primordia, or were born without detectable cardiac abnormalities.^20^ To increase sensitivity for detecting early or subtle phenotypes in our initial screening experiments, we therefore also generated a *PTPN11*^N308D/N308D^ (N308D^Hom^) iPSC line.

In parallel, we also introduced an NSML-associated variant with dominant-negative effects on RAS-MAPK signaling, *PTPN11^Y279C/+^* (Y279C), which is associated with both PVS and hypertrophic cardiomyopathy.^27^ Use of isogenic parental cells allowed us to assign the phenotypic differences we observed across our lines directly to the effects of the pathogenic alleles, rather than variation in genetic background or unknown modifiers. All edits were confirmed with Sanger sequencing (Supplemental Figure 1). Because these edited lines were generated at different times, initial experiments were conducted with the S257L, N308D^Hom^ and Y279C iPSC lines, with N308D^Het^ incorporated in subsequent analyses once it became available.

### Aberrant endocardial differentiation and dysregulated EndMT to fibrosa VICs in NS iPSCs

We previously demonstrated that iPSCs can be successfully driven towards a mesoderm-derived endocardial fate with the GSK-3 inhibitor/WNT activator CHIR-99021 (CHIR), followed by BMP10 and bFGF treatments.^25^ Applying this protocol to our unedited parental line (Ctrl) and CRISPR-edited NS/NSML lines, we were able to generate endocardial cells by Day 14 and, following EndMT, VICs by Day 26 (Figure 1A). By Day 4 of differentiation, we observed an increase in CD13^+^/CD56^+^ cardiogenic mesoderm in the NS lines, but not in the NSML Y279C iPSCs (Supplemental Figure 2A-B). By Day 14, all three mutant lines exhibited a marked increase in the CD31^+^ endocardial cell population relative to Ctrl, with the N308D^Hom^ line producing the most (Figure 1B-D).

**Figure 1:**
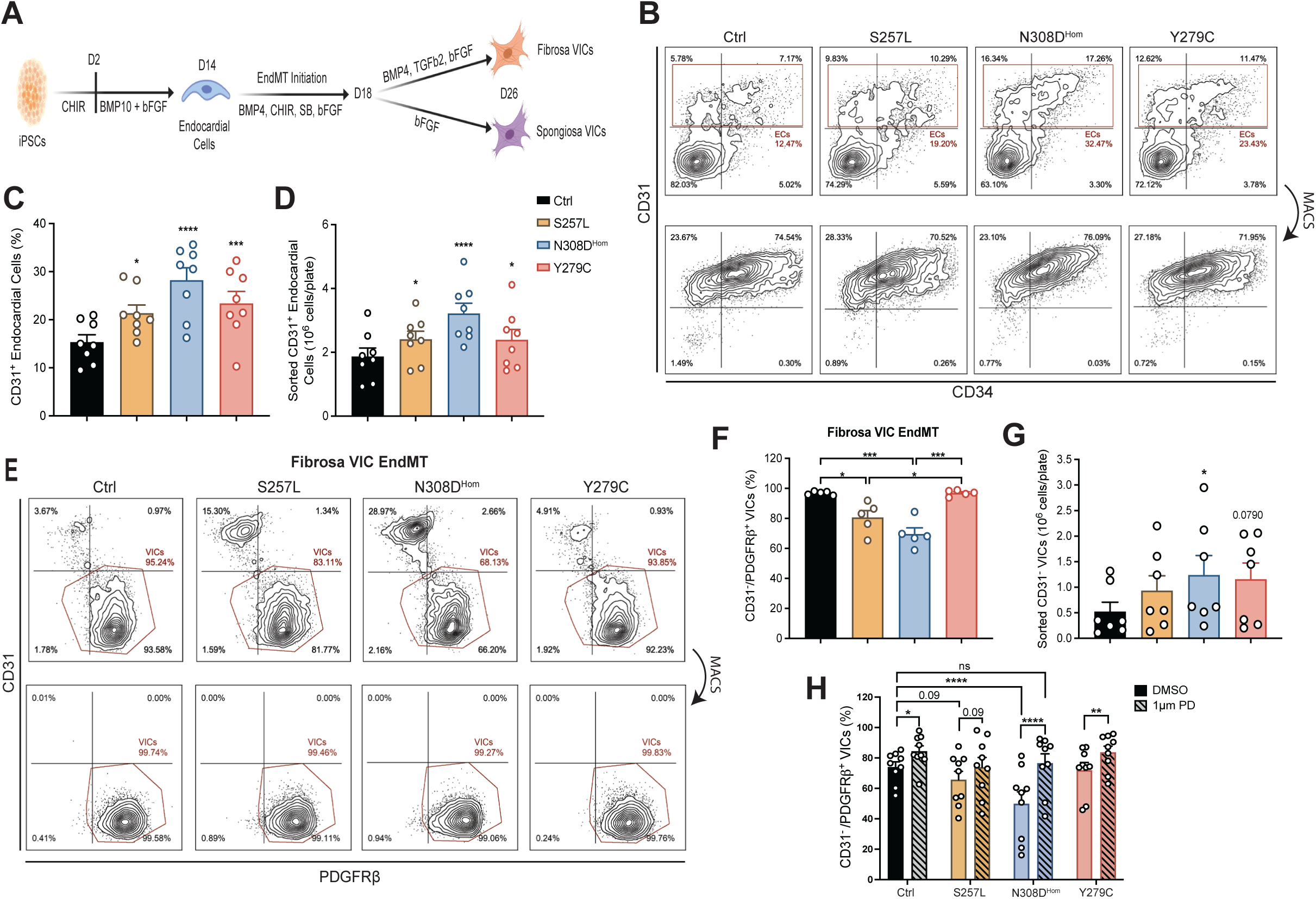
Aberrant differentiation of NS-iPSC lines. **(A)** Schematic of directed differentiation protocol used to generate fibrosa or spongiosa VICs from iPSCs. **(B)** Representative fluorescence-activated cell sorting (FACS) plots of CD31^+^ endocardial-directed cells on Day 14 and enrichment of this population by CD31-directed magnetic-activated cell sorting (MACS). **(C)** Quantification of CD31^+^ endocardial differentiation efficiency on Day 14 prior to MACS. N = 8; *p < 0.05; ***p < 0.001; ****p < 0.0001 by RM one-way ANOVA with Dunnett’s multiple comparisons test, error bars ± SEM. **(D)** Total number of endocardial cells obtained per plate of differentiated iPSCs following CD31-directed MACS. N = 8; *p < 0.05; ****p < 0.0001 by RM one-way ANOVA with Dunnett’s multiple comparisons test, error bars ± SEM. **(E)** Representative FACS plots depicting fibrosa EndMT efficiency on Day 26, measured by generation of CD31^-^/PDGFRβ^+^ VICs, which were subsequently enriched via negative selection with CD31-directed MACS. **(F)** Quantification of fibrosa VIC EndMT efficiency on Day 26 prior to MACS. N = 5; *p < 0.05; ***p < 0.001 by RM one-way ANOVA with Dunnett’s multiple comparison test, error bars ± SEM. **(G)** Total number of fibrosa VICs obtained per plate of differentiated iPSCs following MACS. N = 7; *p < 0.05 by RM one-way ANOVA with Dunnett’s multiple comparisons test, error bars ± SEM. **(H)** Treatment with MEK inhibitor (PD-0325901; 1 µM) during Days 24-26 rescues fibrosa VIC EndMT defect in NS iPSCs. N = 9; *p < 0.05; **p < 0.01; ****p < 0.0001 by two-way ANOVA with Tukey’s multiple comparison test, error bars ± SEM.

To verify that these cells represent *bona fide* endocardial rather than vascular endothelial cells, we generated CD31^+^/CD34^+^ endothelial cells using a VEGF-directed protocol and compared their transcriptional identities (Supplemental Figure 2C-E). RT-qPCR confirmed robust induction of endocardial markers—including *NKX2.5, GATA4, GATA5, ISL1, NRG1, NPR3, HAPLN1, MEIS2,* and *TMEM100*—in cells differentiated with CHIR/BMP10/bFGF. Notably, the expression of these endocardial markers was not significantly altered between our mutant lines and Ctrl, suggesting that endocardial identity is unaltered.

To generate VICs, we next induced EndMT in our endocardial population to generate VICs with a two-stage valve differentiation protocol, including an initial priming phase followed by EndMT in the presence of BMP4, TGFβ2, and bFGF (fibrosa-biased), or only with bFGF (spongiosa-biased) (Figure 1A; detailed methodology is described in our companion study^24^). Under the spongiosa protocol, we observed robust EndMT across all of our lines, based on their ability to generate a CD31^-^/PDGFRβ^+^ VIC population at similar efficiencies (Supplemental Figure 2F-G). In contrast, under the fibrosa protocol, both NS lines had reduced EndMT efficiency—with N308D^Hom^ showing an approximate 30% decrease in CD31^-^/PDGFRβ^+^ VICs compared to Ctrl (Figure 1E-F). This defect was not observed in the NSML line, highlighting known differences in the underlying genetics and molecular mechanisms of NS and NSML.^1,2^

Although N308D^Hom^ exhibited reduced fibrosa EndMT efficiency, this line consistently yielded more total VICs per plate of iPSCs than Ctrl or other mutant lines (Figure 1G). To determine whether increased cell proliferation could account for the increased VIC production, we assessed KI67 expression by flow cytometry (Supplemental Figure 2H-I). We found that less than 10% of fibrosa VICs were KI67^+^ and that no significant differences were observed across genotypes, indicating that altered proliferation does not explain the increased N308D^Hom^ fibrosa VIC yield (Supplemental Figure 2I). Given the central role of increased RAS-MAPK signaling in NS—and supported by our companion study demonstrating enhanced ERK phosphorylation in FGF4-stimulated N308D^Hom^ fibrosa VICs^24^—we next examined whether inhibiting MEK during the final phase of fibrosa EndMT could restore differentiation efficiency. Treating our cells with the MEK inhibitor PD-0325901 (PD; mirdametinib) during the final two days of EndMT (Days 24-26), we observed increased EndMT efficiency across all our lines (Figure 1H). Critically, MEK inhibition fully restored the reduced N308D^Hom^ fibrosa EndMT to Ctrl levels (Figure 1H). These results implicate ERK hyperactivation in impaired EndMT, specifically under fibrosa VIC EndMT conditions.

### scRNAseq characterization confirms robust specification of endocardial and valvular lineages

To characterize the transcriptional identities and differentiation trajectories of our iPSC-derived populations, we performed single-cell RNA sequencing (scRNAseq) on MACS-sorted Day 14 endocardial cells and unsorted Day 26 cells following either spongiosa or fibrosa EndMT protocols. After processing, filtering, and batch correction, we obtained 115,036 high-quality cells for subsequent analysis. UMAP visualization revealed 13 distinct cell clusters, including a *SOX2-*expressing pluripotent cell population; endocardial and valve endothelial cell (VEC) clusters expressing *NPR3* and *PECAM1*; a cycling endocardial cell cluster enriched for *MKI67* (encodes KI-67); and multiple VIC clusters marked by the mesenchymal markers *POSTN* and *COL1A1* (Figure 2A-C). As expected, *PECAM1* expression was higher in the VEC clusters (7 and 8), whereas *NPR3* was enriched within the endocardial clusters (0, 1, and 3) (Figure 2C).

**Figure 2:**
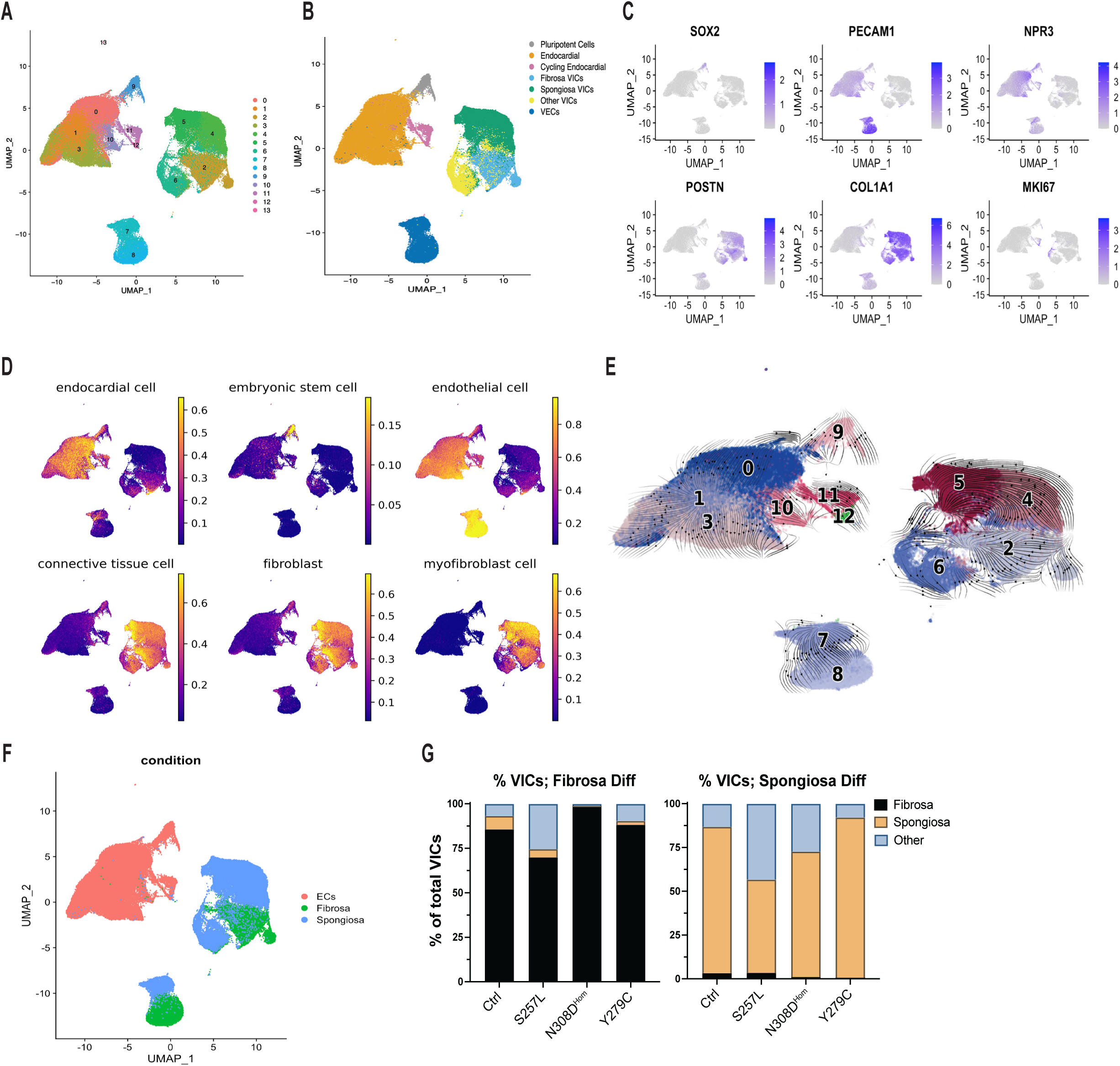
scRNAseq characterization of iPSC differentiation into the cardiac valve lineage. **(A-B)** UMAP visualization of all differentiated cell populations profiled by scRNAseq and cell cluster identities. Numbers denote *Seurat*-defined cell clusters. **(C)** Feature plots showing expression of marker genes for pluripotency (*SOX2)*, endocardial/endothelial cells (*PECAM1)*, endocardial cells (*NPR3)*, valvular interstitial cells (*POSTN, COL1A1)*, and proliferation (*MKI67*). **(D)** *UCDBase*-derived predictions of cluster identities representing absolute cellular phenotypes inferred from deconvolution model fractions. **(E)** RNA velocity streams overlaid on the UMAP, illustrating inferred lineage trajectories. Numbers correspond to the original cluster labels. **(F)** UMAP visualization colored by differentiation protocol, illustrating protocol-defined cell type distribution. **(G)** Relative contributions of cells derived from fibrosa versus spongiosa EndMT protocols within the VIC clusters.

To further validate these identities, we queried *UCDBase*, a deep-learning foundation model for cell type deconvolution.^28^ *UCDBase* accurately identified our *NPR3^+^*cluster as “endocardial cell”, while the *PECAM1^+^* cluster was classified as “endothelial cell” (Figure 2D). Although *UCDBase* was not trained extensively on cardiac valve-specific populations, VIC clusters were annotated as “connective tissue cell,” “fibroblast,” and “myofibroblast cell,” consistent with known VIC phenotypes and functions.

Next, we performed RNA velocity analysis, which leverages spliced and unspliced mRNA abundances to infer cell state transitions and developmental lineage trajectories.^29^ The resulting velocity trajectories indicated movement away from the pluripotent cell cluster toward the endocardial cluster, consistent with progression of iPSCs through a mesoderm-to-endocardial lineage (Figure 2E). Furthermore, we observed divergent velocity trajectories between fibrosa and spongiosa VICs that originate from a common point and end in their respective terminal states, supporting phenotypic differences between these two VIC subtypes. Intriguingly, both VIC protocols also contributed to the “other VICs” cluster (Figure 2F-G). RNA velocity showed that this cluster contributes to both the spongiosa and fibrosa VIC clusters, indicating that this cluster represents a transitional, less mature precursor state that is shared by both VIC subtypes (Figure 2E). Within the VEC cluster, velocity analysis indicated that VECs arising from the spongiosa protocol transition toward the VEC state generated from the fibrosa protocol, implying that the latter may represent a more mature and terminally differentiated phenotype.

Collectively, these transcriptomic data demonstrate that our iPSC-based platform yields *bona fide* endocardial cells, recapitulates the downstream differentiation into the human cardiac valve lineage, and that the introduction of NS- and NSML-associated variants do not alter these cell identities.

### scRNAseq demonstrates enhanced ECM production in NS VICs

Given the aberrant fibrosa EndMT observed in the NS cells (Figure 1E-H), we hypothesized that this VIC subpopulation may play a central role in the development of PVS. Analysis of differentially expressed genes (DEGs) revealed that the N308D^Hom^ fibrosa VICs displayed a particularly robust transcriptional response, with 810 upregulated and 561 downregulated genes compared to the Ctrl fibrosa VICs (Figure 3A). In comparison, the S257L line had 359 upregulated and 268 downregulated DEGs, while the Y279C had 447 upregulated and 264 downregulated DEGs. Gene ontology (GO) analysis for ‘biological process’ terms showed that the top upregulated GO terms across all three mutant fibrosa VIC lines involved terms related to the ECM, such as “extracellular matrix organization” and “collagen fibril organization” (Figure 3B and Supplemental Figure 3A-D). Conversely, downregulated GO terms predominantly involved protein translation and trafficking pathways, such as “cytoplasmic translation” and “protein targeting to ER” (Figure 3C)

**Figure 3:**
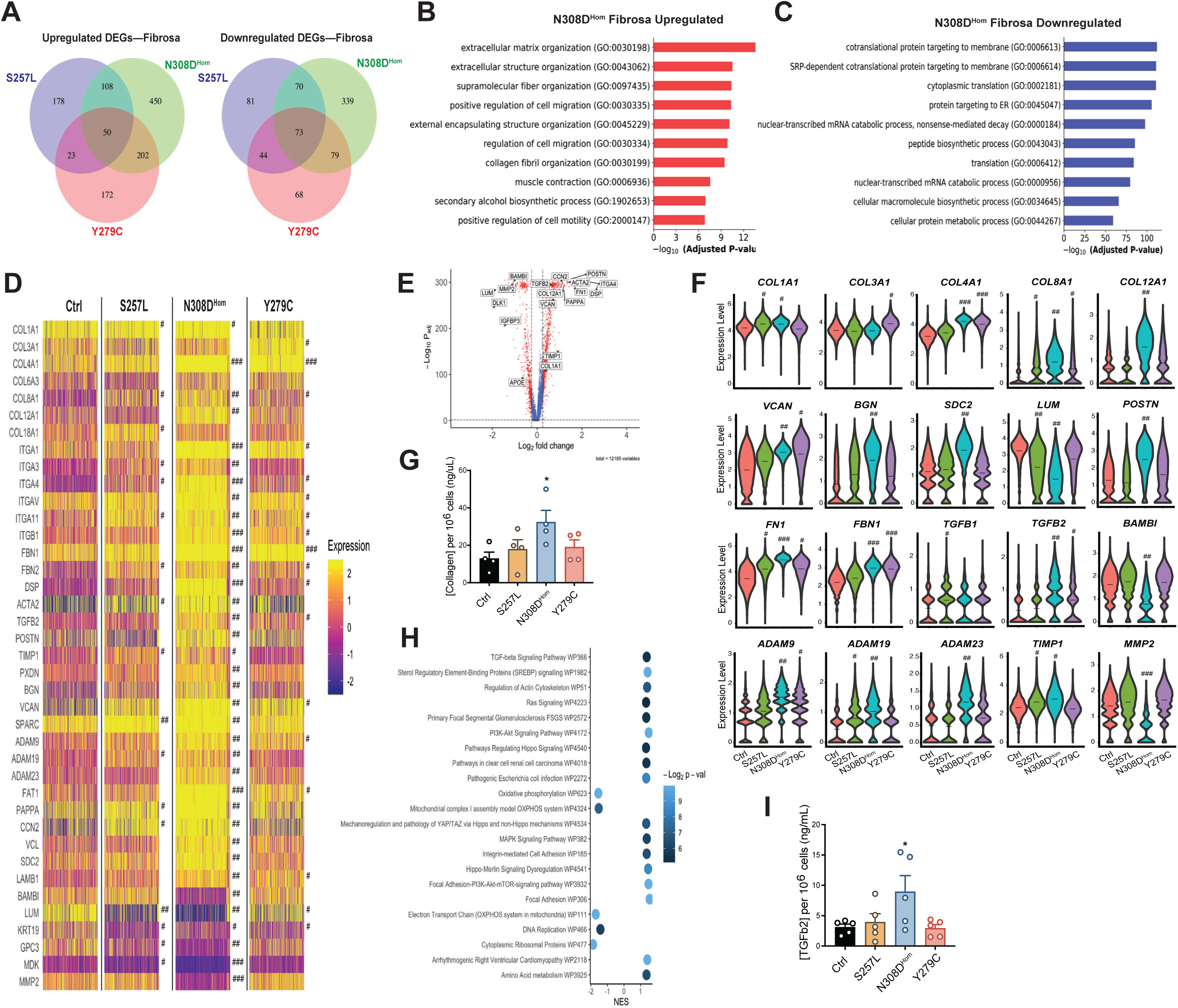
scRNAseq reveals ECM and signaling abnormalities in NS fibrosa VICs. **(A)** Venn diagram showing overlap of upregulated and downregulated DEGs between mutant fibrosa VIC lines. **(B-C)** Gene ontology enrichment analysis of upregulated (B) and downregulated (C) biological processes for N308D^Hom^ fibrosa VICs. **(D)** Heatmap of ECM-related genes that are differentially expressed between Ctrl and NS/NSML cells. Cohen’s *d* effect sizes are indicated as small (#; 0.5 < |*d*| < 1.0), moderate (##; 1.0 < |*d*| < 1.5), or large (###; |*d*| > 1.5). **(E)** Volcano plot of N308D^Hom^ DEGs relative to Ctrl. Dashed lines mark significance thresholds of log_2_ fold change = ± 0.25 and p_adj_ = 0.05. **(F)** Violin plots of ECM-related genes that are differentially expressed between Ctrl and NS/NSML mutants. Dashed lines represent mean expression levels. Cohen’s *d* effect sizes are indicated as small (#), moderate (##), or large (###). **(G)** Total collagen content from fibrosa VIC cultures. N = 4; *p < 0.05 by RM one-way ANOVA with Dunnett’s multiple comparisons test, error bars ± SEM. **(H)** Upregulated and downregulated pathways in N308D^Hom^ fibrosa VICs identified by gene set enrichment analysis. Positive normalized enrichment score (NES) indicates upregulation of the pathway; negative NES indicates downregulation. **(I)** ELISA quantification of TGFβ2 secreted into conditioned media by fibrosa VICs. N = 5, *p < 0.05 by RM one-way ANOVA with Dunnett’s multiple comparisons test, error bars ± SEM.

To further characterize the ECM-associated changes between each of our mutant lines, we focused on genes involved in ECM production, remodeling, and cell-ECM interactions. For each genotype, we also quantified effect sizes relative to Ctrl by calculating Cohen’s *d* score on log-normalized expression values, categorizing effect sizes as small (0.5 < |*d*| < 1.0), moderate (1.0 < |*d*| < 1.5), or large (|*d*| > 1.5). N308D^Hom^ VICs not only showed the broadest upregulation of ECM but also exhibited substantially larger effect sizes compared to the S257L or Y279C (Figure 3D-F). Collagen genes—including *COL1A1, COL4A1, COL8A1,* and *COL12A1—*were strongly upregulated in the N308D^Hom^ fibrosa VICs. Additional matrix genes (*FN1, FBN1, FBN2, POSTN, SPARC)* and proteoglycans (*VCAN, BGN, SDC2*) were similarly elevated. We also observed increased expression of integrins and other cell-matrix interaction genes (*ITGA1, ITGA3, ITGA4, ITGAV, ITGA11, ITGB1, CCN2, ADAM23*). Notably, genes involved in matrix remodeling were dysregulated as well, including upregulation of *TIMP1, ADAM9,* and *ADAM19*, and downregulation of *MMP2*.

We then validated the strong transcriptional signature of enhanced collagen production in N308D^Hom^ cells by quantifying total collagen abundance in Day 26 fibrosa VICs. We detected a nearly two-fold increase in overall collagen production by N308D^Hom^ cells, while both the S257L and Y279C cells were not significantly increased compared to Ctrl (Figure 3G). These findings are consistent with genotype-phenotype correlations in patients, in whom the N308D allele is most strongly associated with PVS development.^8,9^

Because VIC subpopulations may contribute differently to disease, we also examined the spongiosa VICs. This population displayed fewer DEGs overall compared with the fibrosa VICs (Supplemental Figure 4A). In S257L spongiosa VICs, upregulated GO terms were enriched for metabolic processes, such as “glycolytic process” and “pyruvate metabolic process”, whereas N308D^Hom^ spongiosa VICs showed upregulation of pathways related to protein translation and trafficking (Supplemental Figure 4B-E). Notably, both NS spongiosa VIC lines showed downregulation of ECM-related GO terms, supporting the conclusion that the fibrosa VIC lineage is the principal NS cell population driving PVS. In contrast, the NSML Y279C spongiosa VICs displayed the opposite pattern, with upregulation of ECM-related pathways and downregulation of processes associated with cell migration and cell-cell adhesion (Supplemental Figure 4F-G). These findings suggest fundamental mechanistic differences exist between NS-and NSML-associated PVS, with distinct intracellular programs and VIC subpopulations contributing to disease in each condition.

### Dysregulated TGFβ and RAS-MAPK signaling in N308D^Hom^ fibrosa VICs

Because NS and NSML variants affect components of the RAS-MAPK pathway, we next sought to identify putative upstream signaling pathways that could drive the pronounced ECM dysregulation observed in N308D^Hom^ fibrosa VICs. To do this, we performed gene-set enrichment analysis (GSEA) against the *WikiPathways* database. This analysis yielded 22 significantly perturbed pathways (p < 0.05 and FDR < 0.25), including 17 upregulated and five downregulated pathways (Figure 3H). Notably, pathways related to RAS-MAPK, TGFβ, PI3K-AKT, focal adhesion, HIPPO signaling, sterol regulatory element-binding proteins (SREBP), and actin cytoskeleton regulation were upregulated. Downregulated pathways were predominantly metabolic or proliferative processes, including oxidative phosphorylation and DNA replication.

In contrast to the pathway enrichment pattern in N308D^Hom^, GSEA of S257L and Y279C fibrosa VICs revealed notably fewer pathways. In S257L cells, focal adhesion signaling was the only notable upregulated pathway (Supplemental Figure 3E). In Y279C cells, pathways related to focal adhesion, PI3K-AKT, and HIPPO signaling were upregulated, while oxidative phosphorylation was downregulated (Supplemental Figure 3F). When these results were compared against the N308D^Hom^ GSEA signatures, TGFβ and RAS-MAPK signaling pathways emerged as uniquely upregulated in the N308D^Hom^ genotype.

This enrichment of TGFβ signaling was consistent with increased expression of *TGFB2* and concomitant downregulation of *BAMBI*, a negative regulator of TGFβ signaling,^30^ specifically in the N308D^Hom^ cells (Figure 3F). To directly assess ligand production, we measured TGFβ2 levels in conditioned media using ELISA (Figure 3I). Consistent with transcript-level changes, N308D^Hom^ VIC cultures secreted approximately twice as much total TGFβ2 into conditioned media, whereas the S257L and Y279C cultures showed no significant differences.

Collectively, these data indicate that while numerous pathways are perturbed in N308D^Hom^, enrichment of TGFβ and RAS-MAPK signaling appears to be a distinguishing feature of the N308D genotype and likely contributes to the severe phenotype observed clinically in N308D-associated PVS.

### N308D^Het^ largely recapitulates the N308D^Hom^ fibrosa VIC phenotype

Given the strong phenotypic signal observed in the N308D^Hom^ fibrosa VICs, we next sought to determine whether these features were also present in the N308D^Het^ genotype, as NS patients carrying this variant are almost always heterozygous. Establishing concordance between the two genotypes would not only confirm the physiological relevance of our findings but also demonstrate that the observed phenotypic changes are not merely attributable to RAS-MAPK dosage effects arising from differences in zygosity.

Similar to the N308D^Hom^ line, the N308D^Het^ exhibited increased differentiation efficiency toward both mesodermal and endocardial fates, reduced fibrosa EndMT efficiency, and an overall increased VIC yield under the fibrosa differentiation protocol (Figure 4A-H). RT-qPCR analysis further supported our transcriptomic findings, revealing significant upregulation of ECM-associated genes—including *COL1A1, COL8A1, COL12A1,* and *POSTN*—as well as increased expression of *ACTA2* and *ITGA4*, and reduced expression of *BAMBI* (Figure 4I).

**Figure 4:**
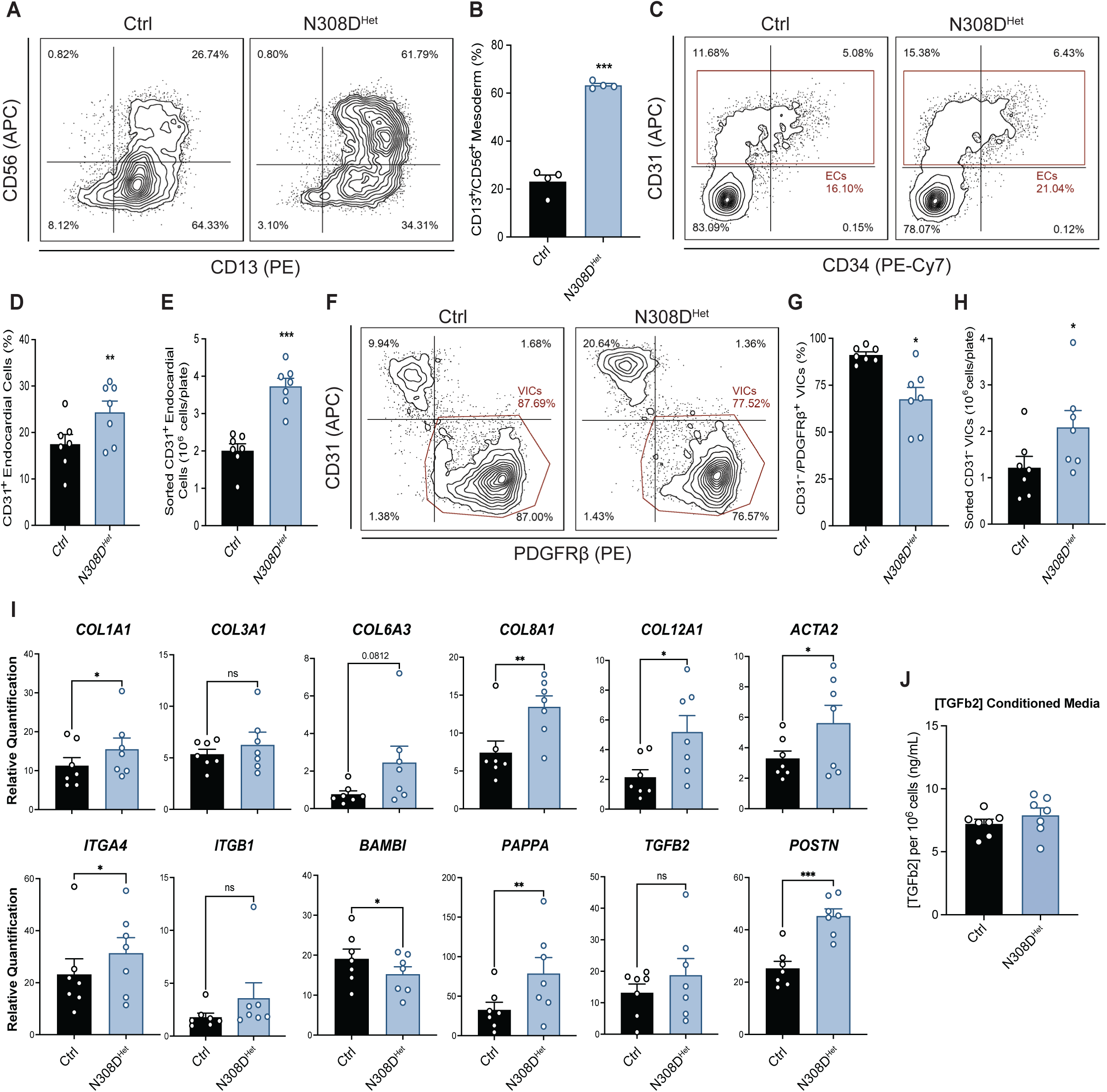
Phenotypic characterization of N308D^Het^ iPSCs. **(A)** Representative FACS plots showing CD13^+^/CD56^+^ mesoderm generation on Day 4 of differentiation. **(B)** Quantification of mesoderm production in N308D^Het^ iPSCs on Day 4. N = 4, ***p < 0.001 by paired t test, error bars ± SEM. **(C)** Representative FACS plots of CD31^+^ endocardial cell generation on Day 14 of differentiation. **(D)** Quantification of endocardial differentiation efficiency in N308D^Het^ cells on Day 14. N = 7; **p < 0.01 by paired t test, error bars ± SEM. **(E)** Total number of endocardial cells obtained per plate of iPSCs differentiated, following CD31-directed MACS. N = 7; ***p < 0.001 by paired t test, error bars ± SEM. **(F)** Representative FACS plots of fibrosa EndMT efficiency on Day 26. **(G)** Quantification of fibrosa EndMT efficiency measured by generation of CD31^-^/PDGFRβ^+^ VICs. N = 7; *p < 0.05 by paired t test, error bars ± SEM. **(H)** Total number of fibrosa VICs obtained per plate of iPSCs differentiated following negative selection by CD31-directed MACS. N = 7, *p < 0.05 by paired t test, error bars ± SEM. **(I)** RT-qPCR of phenotypic markers identified from the scRNAseq dataset in Day 26 Ctrl and N308D^Het^ fibrosa VICs. N = 7; *p < 0.05, **p < 0.01; ***p < 0.001 by paired t test, error bars ± SEM. **(J)** ELISA quantification of TGFβ2 levels in conditioned media from Ctrl and N308D^Het^ fibrosa VICs. N = 7; not significant by paired t test, error bars ± SEM.

Interestingly, unlike the N308D^Hom^ line, the N308D^Het^ cells did not exhibit increased *TGFB2* expression, which was consistent with unchanged TGFβ2 levels in the conditioned media (Figure 4I-J). These data not only suggest that TGFβ2 ligand production is sensitive to gene-dosage effects of the N308D allele, but also that elevated ligand levels may not necessarily be required for dysregulated ECM gene expression in the heterozygous state.

Overall, these results demonstrate that the N308D^Het^ line recapitulates the major phenotypic differences observed in the N308D^Hom^ fibrosa VICs, including aberrant differentiation trajectories and enhanced ECM production, and indicate that the pathogenic consequences of the N308D variant manifest robustly even in the heterozygous genotype.

### Phosphoproteomic analysis reveals global signaling aberrations in N308D^Het^ fibrosa VICs

To define global signaling abnormalities that may underlie the phenotypic changes observed in the N308D^Het^ fibrosa VICs, we performed mass spectrometry-based phosphoproteomic profiling of Day 26 Ctrl and N308D^Het^ cells. After imputation, normalization, and filtering, we quantified 4,499 phosphosites. Principal component analysis (PCA) demonstrated clear genotype separation along PC2, especially after batch correction, with PC2 representing 32.4% of the variance (Figure 5A). This separation indicates widespread, system-level phosphoproteomic differences between Ctrl and N308D^Het^ cells.

**Figure 5:**
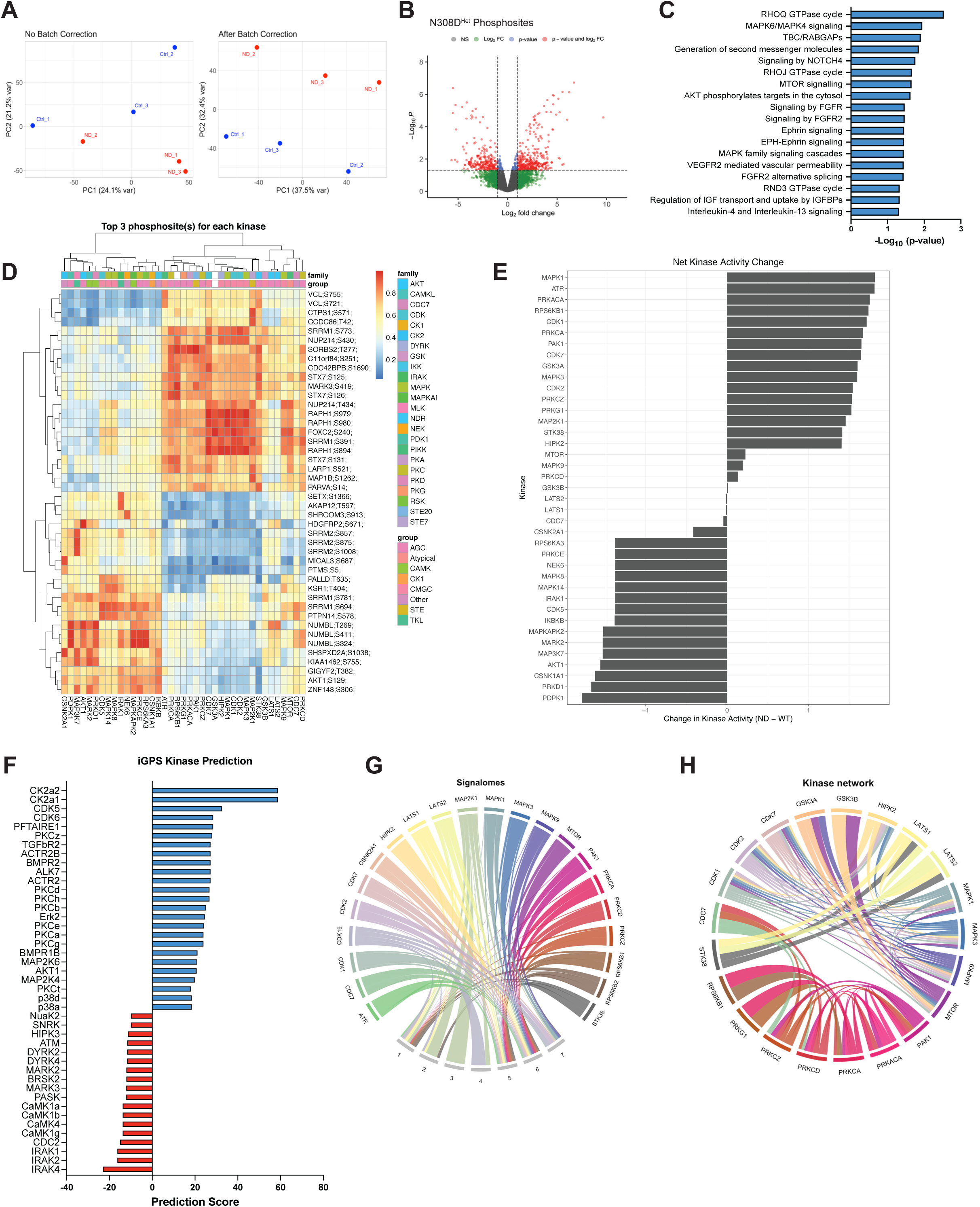
Global phosphoproteomic profiling reveals widespread signaling dysregulation in N308D^Het^ fibrosa VICs. **(A)** Principal component analysis of phosphoproteomic samples before and after batch correction. ND = N308D^Het^ samples **(B)** Volcano plot of differentially regulated phosphosites in N308D^Het^ relative to Ctrl. Dashed lines indicate significance cutoffs of log_2_ fold change = ± 1.0 and p = 0.05. **(C)** Protein-level, rank-based enrichment analysis against the *Reactome* database. **(D)** Clustered heatmap showing the top three predicted substrates for each kinase based on *PhosR* scoring. Higher scores denote stronger predicted kinase-substrate relationships. **(E-F)** Predicted shifts in global kinase activity within N308D^Het^ fibrosa VICs, inferred using *PhosR* (E) and *iGPS* (F). Positive values indicate increased predicted kinase activity; negative values indicate diminished activity. **(G)** Signalome map of upregulated kinase-substrate modules. Branching nodes represent kinases, while stem nodes represent protein modules grouped by shared phosphorylation and regulatory profiles. **(H)** Kinase interaction network illustrating coordinated kinase activity patterns based on substrate relationships.

Differential phosphorylation was then assessed using *limma*,^31^ which identified 63 significant phosphosites (p_adj_ < 0.05). This relatively modest number is consistent with the high biological and technical variability typically observed in phosphoproteomics, particularly in small-sample datasets. To retain sufficient phosphosites for downstream motif enrichment and kinase-inference analyses, we applied a more inclusive threshold (nominal p < 0.05 and |log_2_FC| > 1) which identified 586 altered phosphosites (Figure 5B). We then performed protein-level rank-based enrichment against the *Reactome* database, which revealed upregulation of multiple pathways, including MAPK, AKT-mTOR, RHO GTPase, NOTCH4, FGF, and VEGF signaling (Figure 5C). These pathway shifts broadly paralleled those observed in our transcriptomic analyses.

To gain phosphosite-level insight into the altered signaling pathways, we applied *PhosR*^32^ to infer kinase-substrate relationships across the significantly altered phosphosites. The top three substrates for each kinase were then plotted as a clustered heatmap, revealing two dominant clusters of kinase activity (Figure 5D). The first cluster contained CMGC kinases—including ERK1/2 (MAPK1/3), JNK2 (MAPK9), and CDKs (CDK1/2/7)—as well as PKC isoforms (PRKCA, PRKCD, PRKCZ) and HIPPO kinases (LATS1/2). The second cluster featured alternative MAPKs (JNK1/MAPK8, p38α/MAPK14), CK2 kinases (CSNK1A1, CSNK2A1), PRKCE, NEK6, CDK5, and MARK2. Notably, TAK1/MAP3K7 was identified in this group, which places a key downstream target of noncanonical TGFβ signaling among the kinases with altered activity.^33,34^

Next, we assessed the inferred kinase activity scores in *PhosR* to determine the activity of individual kinases (Figure 5E). This identified robust activation of canonical RAS-MAPK components, including MEK1 (MAP2K1) and ERK1/2 (MAPK1/2). PKC isoforms (PRKCA, PRKCZ) and PKA (PRKACA) also demonstrated increased activity. In contrast, stress-activated kinases, including TAK1/MAP3K7, JNK1/MAPK8, p38α/MAPK14, MAPKAPK2, and PRKD1, had diminished inferred activity. Intriguingly, the AKT-mTOR pathway demonstrated a mixed pattern—activity of mTOR and RPS6KB1 was increased, while AKT1, RPS6KA3, and PDPK1 were decreased—suggesting a shift in pathway activity rather than uniform activation or repression.

Because kinase-substrate inference approaches can differ, we cross-validated these findings with a second kinase-substrate prediction tool, *iGPS*^35^ (Figure 5F). The *iGPS* kinase activity prediction scores corroborated the increased activity of ERK2 and PKC. However, they also suggested that the activity of CK2 kinases, CDK5, and p38α/δ was increased, contrasting with *PhosR* predictions. Importantly, *iGPS* also predicted enhanced activity of TGFbR2, ACTR2B, and BMPR2, indicating activation across the TGFβ receptor superfamily. These predictions align with transcriptional signatures of enhanced TGFβ pathway engagement in our scRNAseq data.

To further resolve kinase-substrate structure, we constructed signalome modules from upregulated phosphosites and identified seven substrate modules (Figure 5G). One module (Module 3) was exclusively regulated by MEK1/MAP2K1, with MEK1 also exerting significant regulation on Modules 2 and 4, highlighting its central role in the phosphoproteome alterations. In addition, CDK19 emerged as a major regulator of Module 4. The remaining substrate modules displayed mixed regulation, with multiple kinases acting on these substrate groups. These module-level relationships were then used to infer a regulatory kinase network (Figure 5H). The resulting kinase map revealed interconnected networks that connect ERK1/2, JNK2, and mTOR pathways to one another. Additionally, we found that this network interacts with CDK1/2/7 and HIPK2. Separately, there is a network between mTOR and GSK3A/B. Notably, the PKC network appears to be distinct from the ERK1/2 network, involving PKC, PKA/PRKACA, PAK1, RPS6K, and PRKG1, with additional interaction from CDC7.

Together, these phosphoproteomic analyses reveal widespread kinase dysfunction in N308D^Het^ fibrosa VICs. The strongest and most consistent signals reflected enhanced RAS-MAPK and PKC activity, accompanied by complex shifts in the activity of stress-activated MAPKs (JNK, p38, TAK1) and AKT-mTOR components. Predictions of increased TGFβ superfamily receptor activity further support a model in which multiple dysregulated kinase networks converge, likely downstream of TGFβ signaling, leading to altered cytoskeletal dynamics, ECM regulation, and response to growth factors.

### Cell-intrinsic defects downstream of TGFβ signaling drives the ECM phenotype in N308D^Het^ fibrosa VICs

Our transcriptomic and phosphoproteomic data implicate enhanced TGFβ signaling in NS fibrosa VICs and that the integration of downstream pathways is aberrant. Additionally, we observed differences in TGFβ2 ligand expression between the N308D^Hom^ and N308D^Het^ fibrosa VICs (Figure 3F-I and Figure 4I-J). To test whether the increased ECM production was due to the abnormal response and integration of TGFβ signaling pathways, rather than the effects of increased TGFβ ligand production, we cultured Day 26 N308D^Het^ fibrosa VICs for three additional days in either the absence or presence of exogenous TGFβ2 (0.3 ng/mL). Following this three-day period, we found that the expression of *COL1A1, COL6A3,* and *COL12A1* was significantly increased in our TGFβ2-treated N308D^Het^ cells compared to TGFβ2-treated Ctrl cells (Figure 6A). Additionally, the expression pattern of *COL1A1, COL6A3,* and *COL12A1* demonstrated that its enhanced expression was dependent on the presence of TGFβ2, as no significant differences were observed between Ctrl and N308D^Het^ cells in the absence of TGFβ2. Importantly, the concentration of TGFβ2 used in these experiments was sufficiently low such that the Ctrl fibrosa VICs failed to exhibit a response. As the N308D^Het^ fibrosa VICs did not exhibit increased production of TGFβ2 ligand (Figure 4I-J), these data implicate NS alleles in driving TGFβ hypersensitivity and a TGFβ2-dependent upregulation of ECM genes.

**Figure 6:**
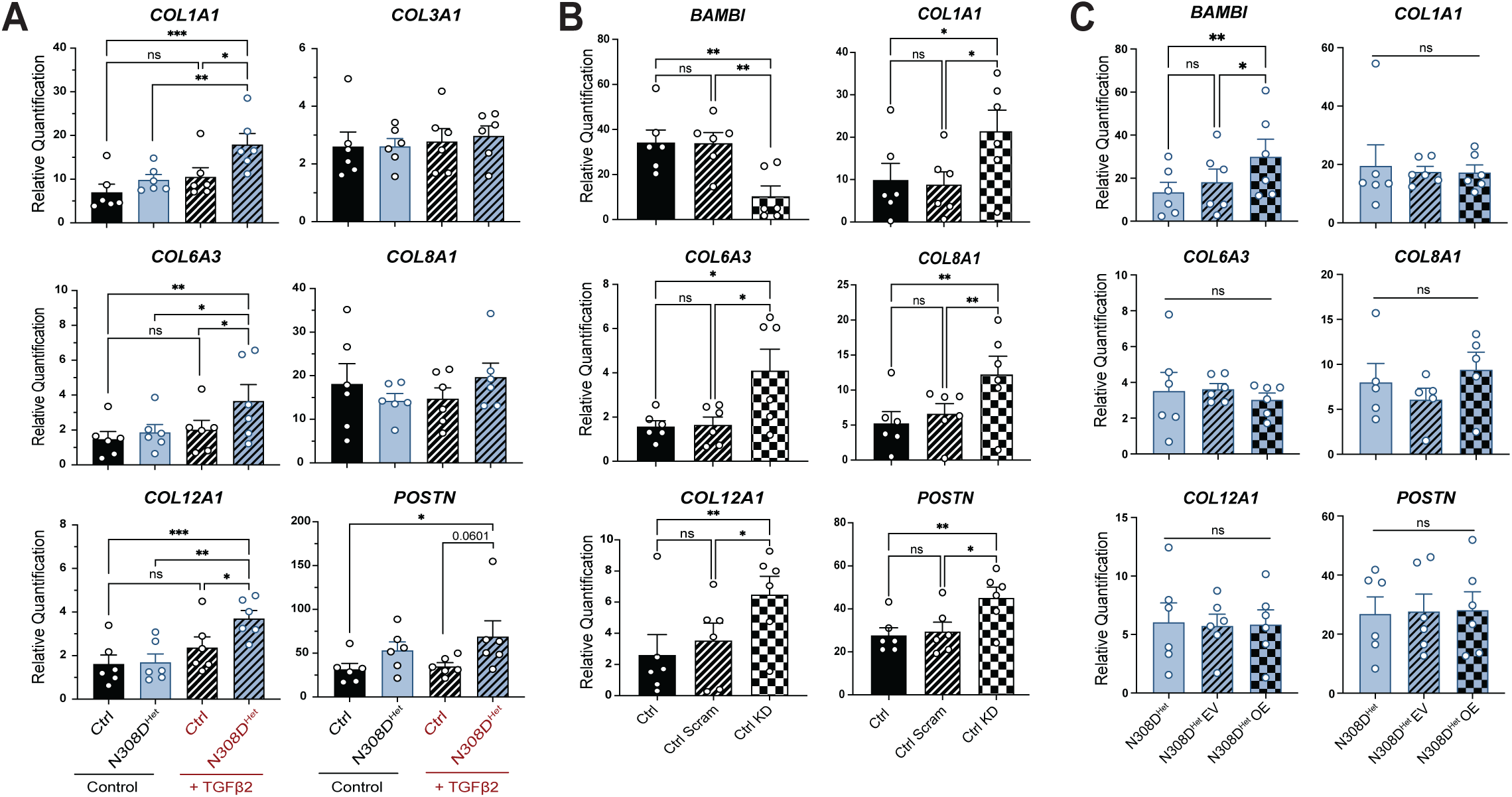
BAMBI-independent hypersensitivity to TGFβ signaling promotes the ECM phenotype in N308D^Het^ fibrosa VICs. **(A)** Enhanced ECM gene expression response in N308D^Het^ fibrosa VICs following TGFβ2 stimulation. N = 6; *p < 0.05, **p < 0.01; ***p < 0.001 by RM one-way ANOVA with Tukey’s multiple comparisons test, error bars ± SEM. **(B)** RT-qPCR of ECM genes following *BAMBI* knockdown in Ctrl fibrosa VICs during EndMT. Scram = scrambled siRNA; KD = knockdown. N = 6; *p < 0.05, ** p < 0.01 by RM one-way ANOVA with Tukey’s multiple comparisons test, error bars ± SEM. **(C)** RT-qPCR of ECM genes following *BAMBI* overexpression in N308D^Het^ fibrosa VICs during EndMT. EV = empty vector; OE = overexpression. N = 6; *p < 0.05, ** p < 0.01 by RM one-way ANOVA with Tukey’s multiple comparisons test, error bars ± SEM.

One potential explanation for these effects is the downregulation of *BAMBI* that we observed in both our N308D^Hom^ and N308D^Het^ fibrosa VICs (Figure 3F and Figure 4I). As a pseudoreceptor of the TGFβ superfamily, BAMBI exhibits an inhibitory effect on TGFβ, BMP, and activin signaling.^30^ To test whether the decreased expression of *BAMBI* could drive the enhanced production of ECM in our cells, we performed siRNA knockdown of *BAMBI* in Ctrl cells throughout fibrosa VIC EndMT (Figure 6B). As expected, transfection of Ctrl cells with *BAMBI* siRNA significantly reduced *BAMBI* expression and resulted in the concomitant upregulation of ECM genes, including *COL1A1, COL6A3, COL8A1, COL12A1,* and *POSTN*.

Next, we asked whether increasing *BAMBI* expression in our N308D^Het^ fibrosa VICs would rescue the ECM phenotype. Transfecting our cells with a *BAMBI* overexpression plasmid throughout fibrosa VIC EndMT, we were able to detect increased *BAMBI* expression by Day 26 (Figure 6C). However, despite this increase in *BAMBI*, we did not observe any significant changes in *COL1A1, COL6A3, COL8A1, COL12A1,* or *POSTN* expression.

These data suggest that the integration and interpretation of signals downstream of TGFβ are abnormal in N308D^Het^ fibrosa VICs, resulting in increased synthesis of ECM proteins. Furthermore, we demonstrate that although diminished *BAMBI* expression is sufficient for increasing ECM production in Ctrl cells, restoring its expression is unable to rescue the pathological ECM program in N308D^Het^ fibrosa VICs. As such, BAMBI-independent pathways likely exist to maintain excessive ECM gene expression.

### Noonan syndrome patient pulmonary valves exhibit disordered and increased deposition of ECM

Our *in vitro* modeling of NS-associated PVS suggested that excessive and dysregulated ECM production is a central driver of valvular stenosis. To validate these findings in human tissues, we examined pulmonary valves from two infants with NS (ages 4 and 5 months) undergoing valvotomy surgery. These patients were found to have *PTPN11* N308T (NS-PT1) and *RAF1* P261A (NS-PT2) variants (Supplemental Figure 5A). For comparison, we obtained three normal, age-matched pulmonary valves from patients without NS. Since human heart valves mature and remodel their ECM throughout early life and into adolescence,^14^ it is critical that comparisons between valve tissues are made using specimens similar in age.

Grossly, normal pulmonary valves appeared thin and smooth, while the NS valves were dysmorphic and overtly thickened. On histological examination with hematoxylin and eosin (H&E) staining, we found that the NS valves were not only larger and contained more ECM but also appeared to be hypercellular—most prominently in NS-PT2 (Figure 7A; Supplemental Figure 5B). Notably, we observed a stark difference in the morphology of the nuclei, with NS valves displaying elongated, spindle-shaped nuclei compared to the rounded nuclei typically observed in the control valves, suggestive of ongoing VIC activation or stress in NS valves (Supplemental Figure 5B).

**Figure 7:**
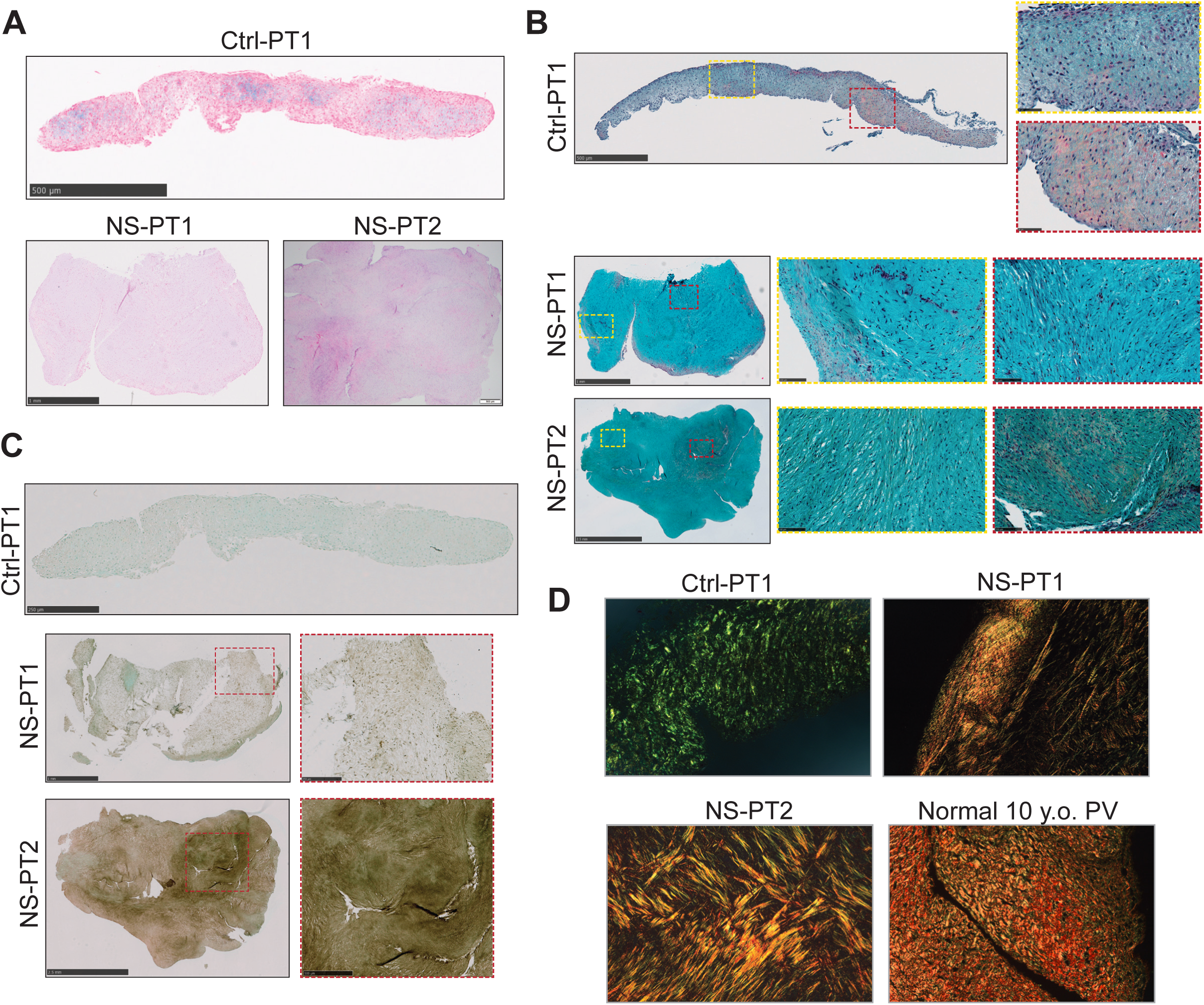
Histopathological analysis reveals ECM disorganization in NS stenotic pulmonary valves. **(A)** Representative hematoxylin and eosin (H&E) staining of normal (Ctrl-PT1) and NS-associated stenotic pulmonary valves (NS-PT1, NS-PT2). **(B)** Movat pentachrome staining of pulmonary valves, highlighting collagen/reticular fibers (yellow), elastic fibers (black), ground substance/mucin (blue), and fibrin/muscle (red). Dashed rectangles indicate regions shown at higher magnification. **(C)** Immunohistochemical staining for versican in pulmonary valve sections. Brown staining denotes versican expression; counterstain performed with toluidine blue. **(D)** Picrosirius red staining visualized microscopically under polarized light to differentiate fibrillar collagen subtypes and fibril organization. Type III collagen appears green, while type I collagen exhibits yellow-orange birefringence.

To assess ECM composition, we performed Movat pentachrome staining, which distinguishes between collagen/reticular fibers (yellow), elastic fibers (black), ground substance/mucin (blue), and fibrin/muscle (red).^36^ Control valves demonstrated the expected spectrum of ECM proteins, with a predominance of ground substance and areas containing collagen and fibrin fibers (Figure 7B; Supplemental Figure 5C). In contrast, the NS valves exhibited intense blue staining, with only focal yellow and red regions. This bright blue staining pattern suggests substantial amounts of ground substance, which includes proteoglycans and glycoproteins. ECM fibers were also visibly disorganized throughout the NS valves, with loss of the typical lamellar alignment found in the controls. Motivated by these findings, we then stained for the presence of versican, a proteoglycan that was upregulated in our transcriptomic data and contributes to myxomatous valve degeneration^37,38^ (Figure 7C; Supplemental Figure 5D). Versican expression was minimal in the control valves but was substantially increased in the NS valves, particularly in NS-PT2.

To further characterize collagen fiber architecture within the ECM, we performed picrosirius red staining followed by polarized light microscopy, which distinguishes between type I (yellow-orange birefringence) and type III (green birefringence) collagen.^39^ In the control valves, we detected type III collagen throughout the valve, with only small punctate areas of type I collagen colocalization, consistent with the nascent ECM of the early postnatal valve leaflet (Figure 7D). In contrast, NS valves showed a robust type I collagen signal that was closely interwoven with type III collagen. Additionally, the collagen fibrils in NS valves exhibited a cross-hatching pattern indicative of collagen fiber disorganization. As an additional control, we also obtained a normal pulmonary valve from a 10-year-old patient, which demonstrated significant maturation of the matrix, consisting of increased type I collagen expression that colocalized with type III collagen (Figure 7D). Notably, the collagen fibrils in this valve were well-organized and provide a stark contrast to that of the NS valves.

Collectively, these histological analyses demonstrate widespread ECM abnormalities in NS patient valves, including excessive ground substance deposition, a shift from type III to type I collagen, and profound disruption of ECM architecture. These findings strongly corroborate our *in vitro* model and are consistent with a TGFβ-dependent mechanism of ECM dysregulation underlying NS-associated PVS.

## Discussion

Pulmonary valve stenosis is one of the most common and clinically important manifestations of NS, yet its developmental and molecular underpinnings are incompletely defined. In this study, we used a human iPSC-derived valve differentiation platform that permits specification of fibrosa and spongiosa VIC subtypes.^24,25,40^ Using CRISPR-edited NS and NSML iPSCs, we examined how these pathogenic variants alter human valve development and signaling. By integrating cell lineage phenotyping, single-cell transcriptomics, phosphoproteomics, *in vitro* assays, and histological analysis of stenotic pulmonary valves from infants with NS, we identify a central mechanism in which NS variants render fibrosa VICs hypersensitive to TGFβ signaling. This heightened TGFβ responsiveness is associated with activation of non-canonical TGFβ pathways and occurs through a BAMBI-independent mechanism. The net result is excessive and disorganized ECM deposition that characterizes NS-associated PVS.

### Abnormal valvular differentiation underpins development of NS-associated PVS

Across each of our pathogenic alleles, we observed aberrant development throughout differentiation, including enhanced generation of mesodermal and endocardial populations. Additionally, N308D variants also produced more fibrosa VICs overall. These findings parallel observations in NS mouse models, where enlarged endocardial cushion and valve primordia have been reported during fetal development.^19,20^ Interestingly, although overall fibrosa VIC numbers were increased, EndMT efficiency toward fibrosa VICs was reduced and occurred in an ERK-dependent mechanism. We did not detect differences in VIC proliferation, suggesting that increased production of endocardial precursors is a major contributor to the enlarged fibrosa VIC population. Together, these findings indicate that NS variants perturb early developmental programs governing valvulogenesis and suggest the presence of additional cell-autonomous mechanisms that contribute to NS-associated PVS.

### NS fibrosa VICs are pathologically primed for TGFβ activation

TGFβ signaling, particularly via TGFβ2, is essential for normal valvulogenesis, regulating EndMT, ECM production, and matrix remodeling.^41,42^ TGFβ also plays a central role in fibroblast biology, mediating activation, proliferation, and ECM production, while excessive TGFβ signaling is a hallmark of pathological organ and tissue fibrosis.^43^ The contribution of TGFβ signaling to NS-associated PVS, however, has not been previously defined.

Several lines of evidence from this study support enhanced and dysregulated TGFβ signaling as a central driver of NS-associated PVS. Our transcriptomic data revealed widespread disruption of ECM gene expression in N308D^Hom^ fibrosa VICs, including upregulation of type I collagen (*COL1A1)*, network-forming collagens (*COL4A1, COL8A1)*, fibril-associated collagens with interrupted triple-helices (FACITs) (*COL12A1)*, and proteoglycans (*VCAN, BGN, SDC2)*. We also observed increased activation of NS fibrosa VICs, marked by elevated *ACTA2* expression, indicating upstream signaling aberrations driving the upregulation of fibrotic gene programs. GSEA highlighted increased activity of the TGFβ and RAS-MAPK pathways, as well as non-canonical TGFβ pathways such as AKT-mTOR and RHO-GTPase, which regulate the actin cytoskeleton.^44,45^ Additional pro-fibrotic pathways that intersect with TGFβ signaling, including integrin-focal adhesion and HIPPO pathways, were also enriched.^46,47^

Phosphoproteomic profiling corroborated these global signaling disruptions in NS-associated PVS. The network-level shifts in kinase activity indicated robust activation of ERK1/2 and PKC, both of which are implicated in TGFβ-driven production of ECM and fibrotic responses in fibroblasts.^48,49^ The intersection of NS-mediated RAS-MAPK activation with non-canonical TGFβ signaling highlights a plausible feedback mechanism whereby sustained ERK activity enables hyperactivation of TGFβ signaling, which then further potentiates ERK activity. In fact, there is evidence that TGFβ-mediated effects on epithelial-mesenchymal transition and fibroblast activation require ERK1/2 activity, as MEK inhibition attenuates these effects of TGFβ.^50,51^

In addition to ERK and PKC, we detected altered activity in AKT-mTOR kinases and stress-activated MAPKs, including JNK, p38, and TAK1. TAK1 is particularly notable given its key role in mediating non-canonical TGFβ signaling through JNK and p38, and its established involvement in ECM production and fibrosis.^33,34,52^ Consistent with these findings, upstream kinase prediction with *iGPS* identified TGFβ receptor activation as a major driver of phosphosite changes in N308D^Het^ cells. Collectively, these data indicate NS variants create a signaling state in which multiple TGFβ-associated pathways converge to amplify ECM production in fibrosa VICs.

### BAMBI-independent hypersensitivity to TGFβ drives NS-associated PVS

To directly assess whether the N308D^Het^ fibrosa VICs have altered responses to TGFβ ligand, we exposed them to exogenous TGFβ2. These cells exhibited an exaggerated ECM response to the presence of TGFβ2 ligand, despite exposure to the same concentration of ligand as Ctrl cells. Given that TGFβ2 ligand production is not enhanced in N308D^Het^ cells, the response we observe suggests an intrinsic difference in signal interpretation, rather than simply ligand availability.

Reduced expression of *BAMBI*—as observed in our transcriptomic data—provides a plausible mechanism for this altered response. As a pseudoreceptor, BAMBI lacks an intracellular kinase domain and prevents complex formation and activation of TGFβ receptors.^30^ *BAMBI* expression is also induced by TGFβ signaling, highlighting its importance in the negative feedback regulation of TGFβ signaling.^53^ Interestingly, reduced BAMBI levels have also been reported in pediatric congenital aortic valve stenosis tissues and have been proposed to act as a master regulator of the valve collagen interactome.^54^

We therefore tested the effects of modulating *BAMBI* expression during valve differentiation. Our results showed that reducing *BAMBI* expression is sufficient to drive ECM upregulation in a Ctrl background. However, restoring *BAMBI* levels in N308D^Het^ fibrosa VICs did not revert the high-ECM phenotype. This indicates that diminished *BAMBI* is not required for the ECM phenotype observed in N308D^Het^ fibrosa VICs and that additional BAMBI-independent pathways—such as sustained ERK activation—maintain the hypersensitivity to TGFβ signaling and the pathogenic ECM program of NS-associated PVS.

### Patient valve histopathology mirrors the phenotypic consequences observed in NS iPSCs

Histological analysis of stenotic pulmonary valves from two infants with NS revealed pronounced and disorganized ECM accumulation, including increased ground substance, elevated versican deposition, a shift from type III to type I collagen, and widespread collagen fibril disorganization. The increased deposition of type I collagen—which is mechanically stiffer than the more flexible type III collagen—is consistent with studies demonstrating that an increased type I:type III collagen ratio increases tissue stiffness and promotes fibroblast activation.^55,56^ In contrast, myxomatous mitral valve disease features the opposite pattern, with increased type III and reduced type I collagen leading to weakened leaflet ECM architecture and prolapse.^57^ This further highlights the pathogenic role of excess type I collagen in driving increased stiffness and stenosis of NS-associated pulmonary valves.

Our histological findings also closely parallel the transcriptomic signatures observed in N308D^Het^ fibrosa VICs, including upregulation of *COL12A1* and *VCAN*—both of which directly interact with and regulate type I collagen fibril organization^16,58–60^—as well as dysregulation of key ECM-remodeling regulators critical for normal valve development, such as *TIMP1, ADAM9, ADAM19,* and *MMP2.*^61–63^ To the best of our knowledge, these analyses represent the first detailed histopathological characterization of NS-associated PVS and provide strong validation of our iPSC-derived valve cell model.

### Clinical insights into NS-associated PVS

In patients with NS, PVS is often mild to moderate at birth but frequently progresses during the first year of life.^64^ Current management of severe disease relies on balloon valvuloplasty, which is rarely effective and necessitates repeat interventions, or surgical valvectomy, which often requires subsequent valve replacement later in life.^64–66^ These limitations highlight the need for novel pharmacological interventions for these patients. The characteristic postnatal progression suggests that disease pathogenesis is driven not only by defective *in utero* valvulogenesis, but also by intrinsic differences in VIC biology during the critical period of postnatal remodeling. Intriguingly, early evidence suggests that compassionate use of the MEK inhibitor trametinib in patients with severe NS-associated hypertrophic cardiomyopathy can result in echocardiographic improvement of right ventricular outflow tract flow gradients, indicating reduced severity of PVS.^67,68^ Moreover, trametinib was recently administered to an infant with NS due to an autosomal recessive *LZTR1* variant who presented with severe pulmonary stenosis and mild hypertrophic cardiomyopathy; in this case, the peak instantaneous valvar gradients decreased from ∼85 mmHg to < 20 mmHg after nine months of continued treatment (Gelb *et al*., unpublished). These observations suggest that NS-associated PVS is at least partially reversible, that a window of postnatal valve plasticity may exist, and that ERK1/2 signaling is central to disease pathogenesis. They also underscore the need to define the relevant pathways in detail and to develop novel therapies—such as TGFβ inhibition—for patients who are intolerant of, or unresponsive to, MEK inhibition.

### Conclusions and implications

Our findings support a model in which NS variants disrupt early developmental programs and produce a fibrosa VIC population that is pathologically hypersensitive to TGFβ signaling. Through ERK-dependent and BAMBI-independent mechanisms, this sensitized cellular state promotes the pathological expression, deposition, and structural remodeling of ECM proteins in NS-associated PVS. By integrating human iPSC valve differentiation, single-cell transcriptomics, phosphoproteomic profiling, and histopathological analysis of patient valves, we identify dysregulated TGFβ-RAS-MAPK signaling as a central mechanism responsible for NS-associated PVS. More broadly, this work establishes an *in vitro* platform for mechanistic interrogation of human valvular disease and provides a framework for developing targeted strategies beyond MEK inhibition to prevent or reverse NS-associated PVS.

## Acknowledgments

We extend our gratitude to Dr. Nicole Dubois (Icahn School of Medicine at Mount Sinai) and her lab members for fruitful discussions and guidance throughout this project. Additionally, we are eternally grateful for the patients and families who have generously permitted the use of their surgical specimens for research purposes. This work was conducted with the support of the Flow Cytometry CoRE and the Biorepository and Pathology CoRE at the Icahn School of Medicine at Mount Sinai. Computational support was provided in part through the Minerva computational and data resources and staff expertise provided by Scientific Computing and Data at the Icahn School of Medicine at Mount Sinai and supported by the Clinical and Translational Science Awards (CTSA) grant UL1TR004419 from the National Center for Advancing Translational Sciences. This work was also supported in part by the Medical Scientist Training Program (MSTP) at the Icahn School of Medicine at Mount Sinai via the NIH T32 training grant 1T32GM146636. The Sidoli lab gratefully acknowledges for funding the Hevolution Foundation (AFAR), the Einstein-Mount Sinai Diabetes center, the ERC-CFAR center for AIDS research, and the NIH Office of the Director (S10OD030286). Illustrations in this manuscript depicting the differentiation schematics were created using BioRender.com.

---

**Supplemental Figure 1:**
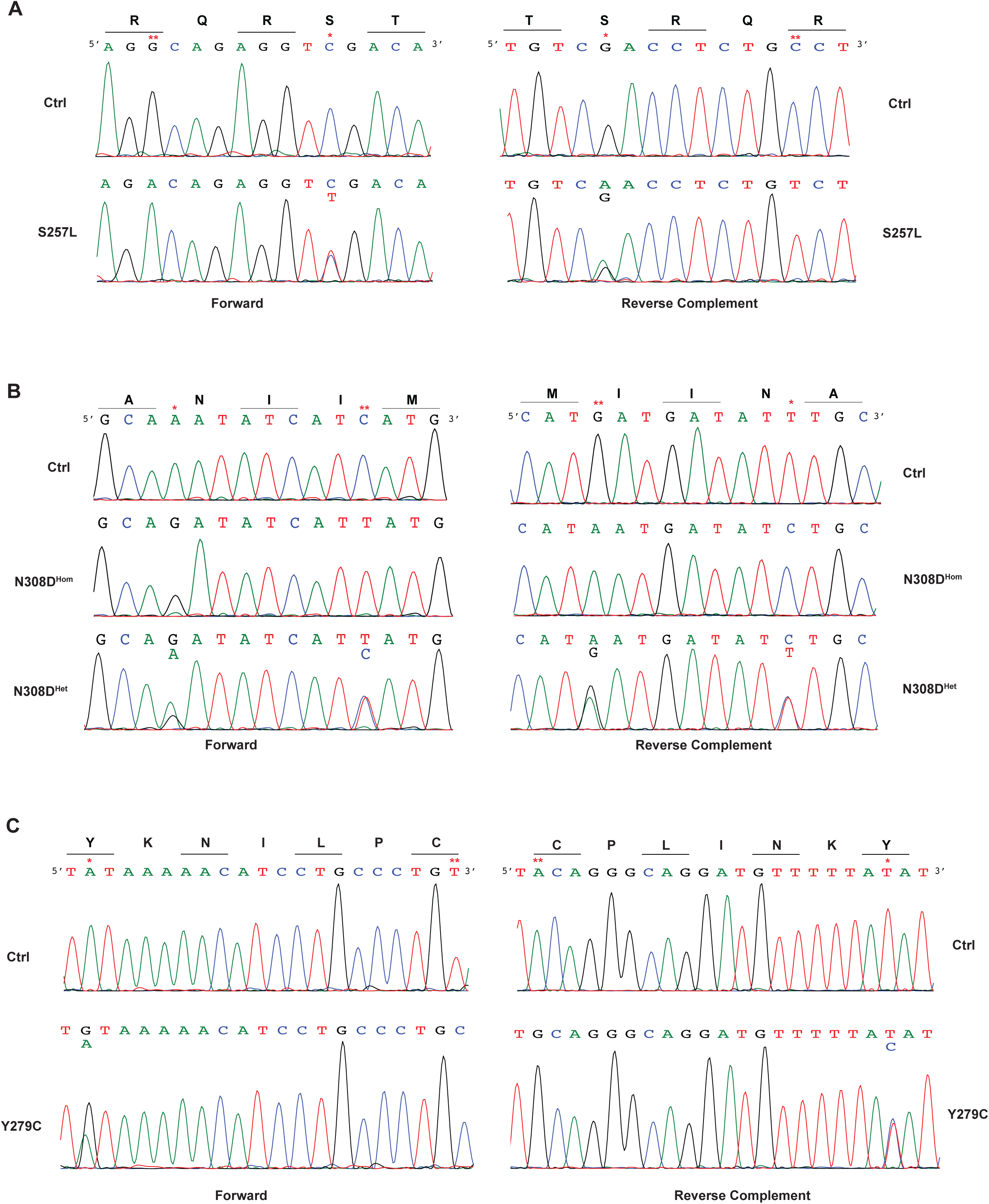
Sanger sequencing of CRISPR-edited NS and NSML iPSC lines.

**Supplemental Figure 2:**
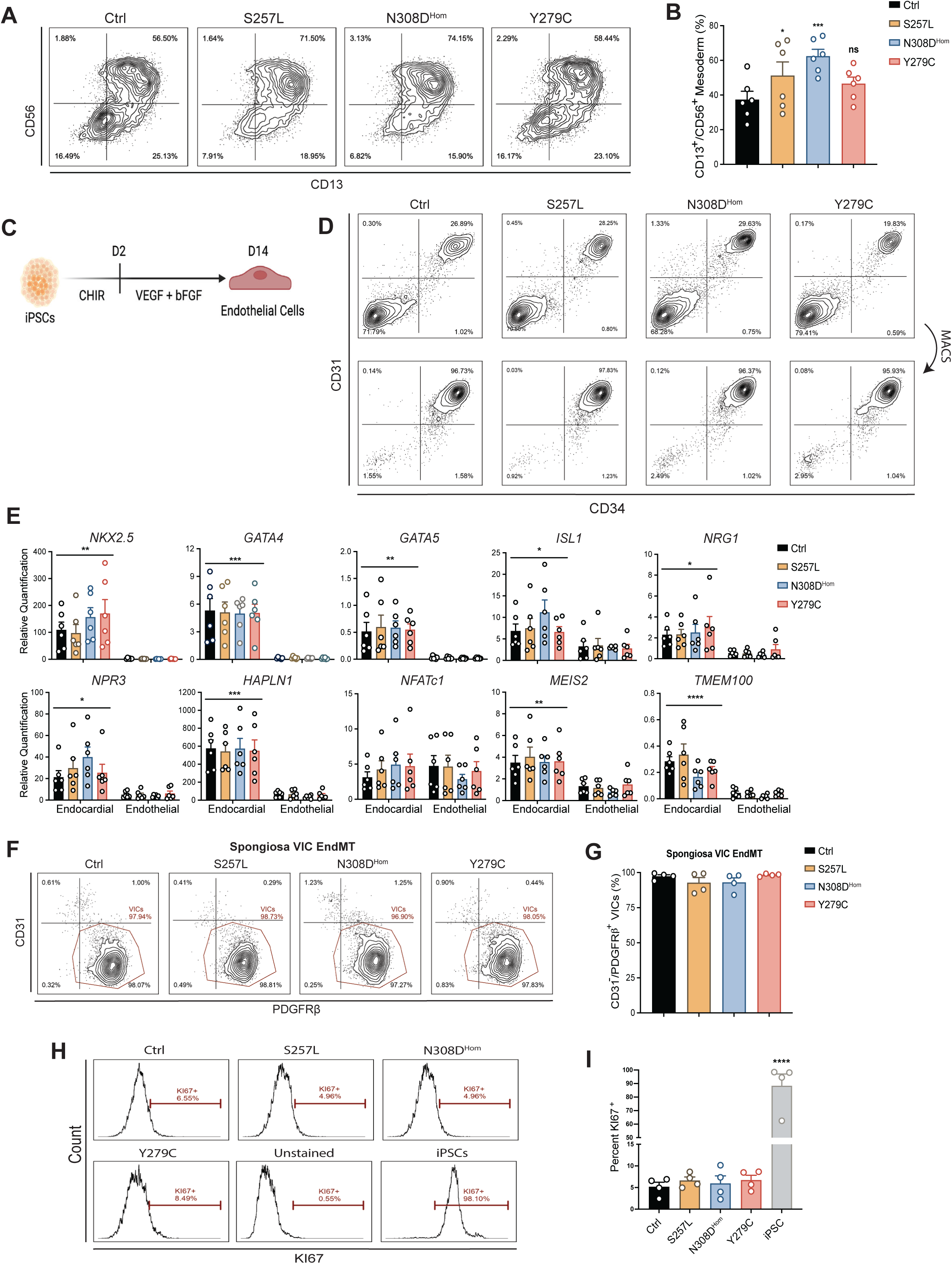
Characterization of NS and NSML iPSCs during differentiation to valvular cells.

**Supplemental Figure 3:**
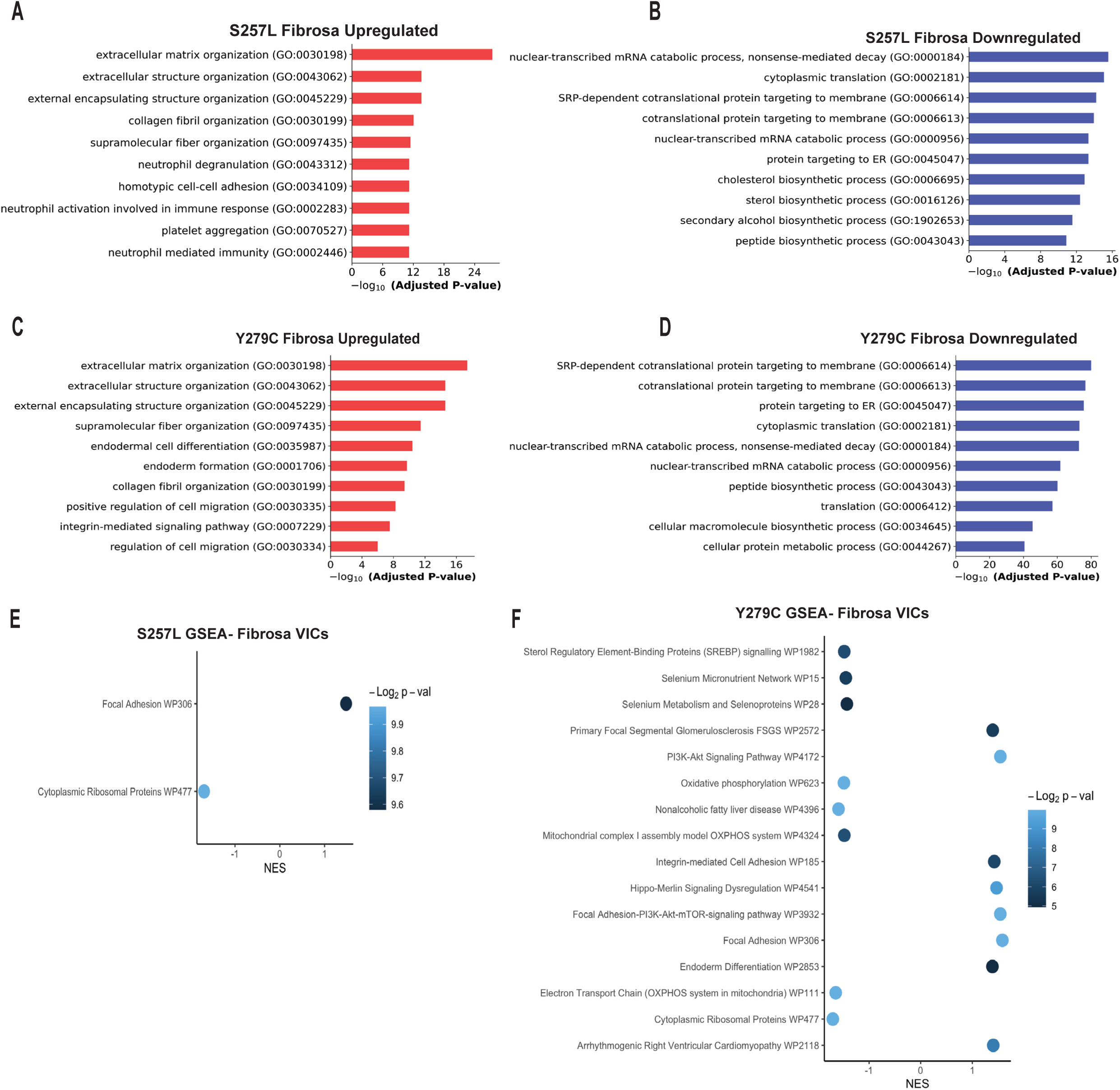
scRNAseq characterization of S257L and Y279C fibrosa VICs.

**Supplemental Figure 4:**
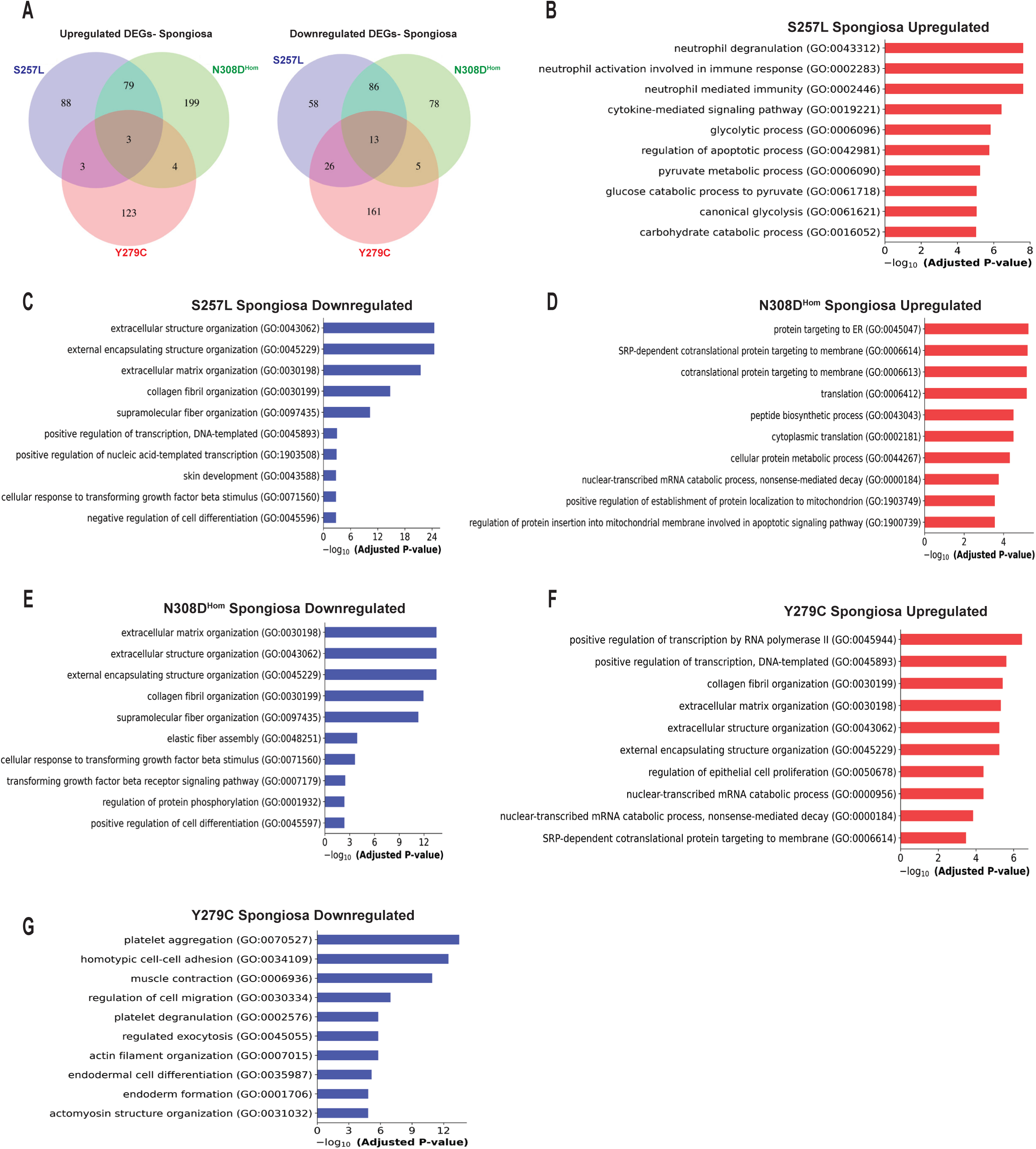
scRNAseq characterization of spongiosa VICs.

**Supplemental Figure 5:**
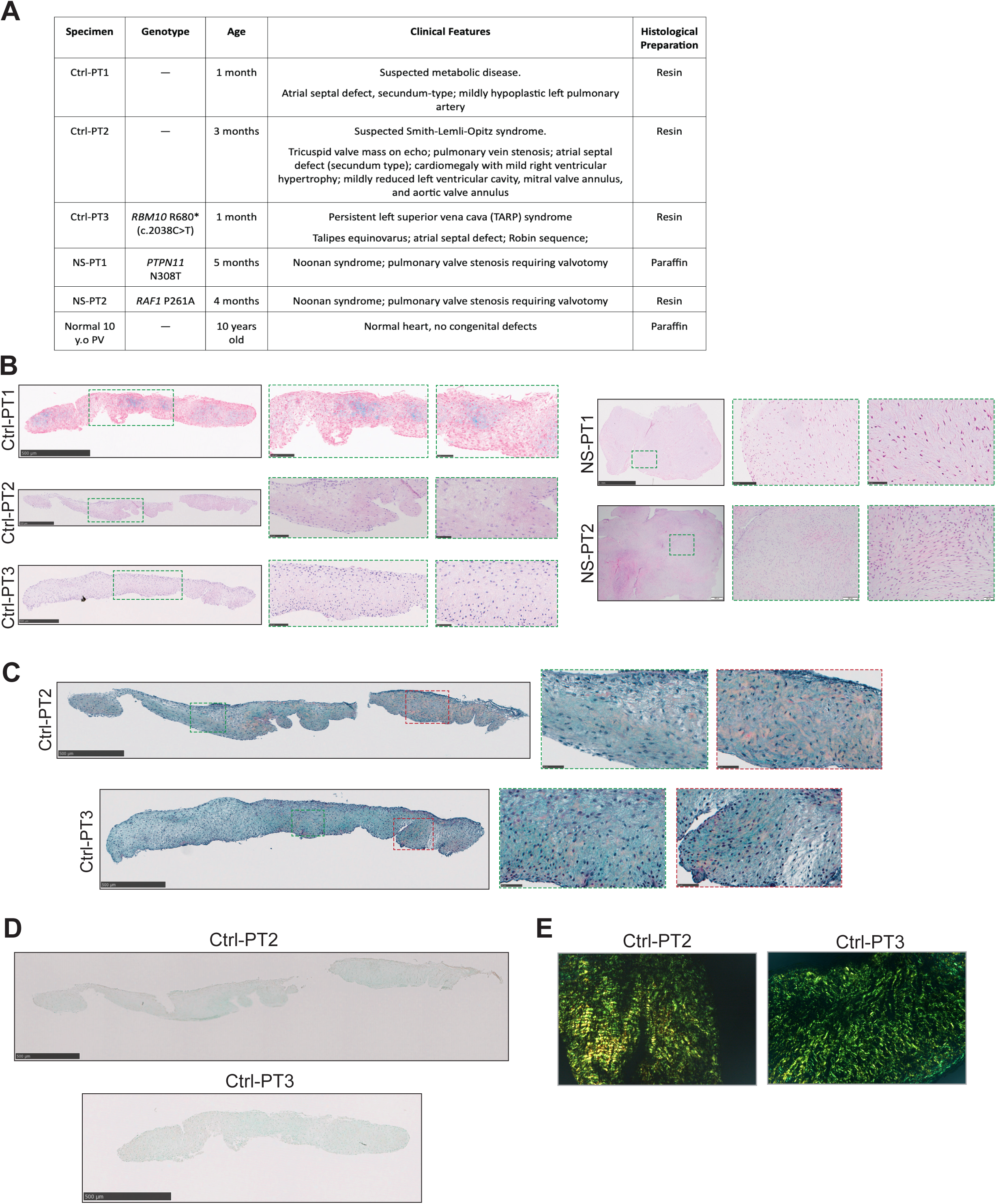
Additional histological characterization of normal and NS-associated stenotic pulmonary valves.

